# Brain virtual histology with X-ray phase-contrast tomography Part I: whole-brain myelin mapping in white-matter injury models

**DOI:** 10.1101/2021.03.24.436852

**Authors:** Matthieu Chourrout, Hugo Rositi, Elodie Ong, Violaine Hubert, Alexandre Paccalet, Louis Foucault, Awen Autret, Barbara Fayard, Cécile Olivier, Radu Bolbos, Françoise Peyrin, Claire Crola-da-Silva, David Meyronet, Olivier Raineteau, Hélène Elleaume, Emmanuel Brun, Fabien Chauveau, Marlène Wiart

**Author notes:** **Corresponding author**: Marlène WIART, Inserm U1060 CARMEN-IRIS, Groupement Hospitalier EST, bâtiment B13, IHU OPERA, 59 boulevard Pinel 69500 BRON -FR, Mail. Co-first authors (Matthieu Chourrout, Hugo Rositi). co-last authors (Fabien Chauveau, Marlène Wiart). **Author contributions** (according to Contributor Role Taxonomy (<u>CRediT</u>)): Conceptualization: MW, FC Data curation: MC, EB, HR, MW, FC Formal analysis: MC, AA Funding acquisition: EB, FP, MW, FC Investigation: HR, EO, VH, AP, CCdS, LF, OR, RB, EB, CO, FP, FC, MW Methodology: EB, HR, MW, FC Project administration: MW, FC Resources: BF, DM Software: AA, BF Supervision: EB, FC, MW Validation: MC, HE, FP, FC, MW Visualization: MC Writing – original draft: MW Writing – review & editing: MC, FP, HE, HR, MW, FC.

## Abstract

White-matter injury leads to severe functional loss in many neurological diseases. Myelin staining on histological samples is the most common technique to investigate white-matter fibers. However, tissue processing and sectioning may affect the reliability of 3D volumetric assessments. The purpose of this study was to propose an approach that enables myelin fibers to be mapped in the whole rodent brain with microscopic resolution and without the need for strenuous staining. With this aim, we coupled inline (propagation-based) X-ray phase-contrast tomography (XPCT) to ethanol-induced brain sample dehydration. We here provide the proof-of-concept that this approach enhances myelinated axons in rodent and human brain tissue. In addition, we demonstrated that white-matter injuries could be detected and quantified with this approach, using three animal models: ischemic stroke, premature birth and multiple sclerosis. Furthermore, in analogy to diffusion tensor imaging (DTI), we retrieved fiber directions and DTI-like diffusion metrics from our XPCT data to quantitatively characterize white-matter microstructure. Finally, we showed that this non-destructive approach was compatible with subsequent complementary brain sample analysis by conventional histology. In-line XPCT might thus become a novel gold-standard for investigating white-matter injury in the intact brain. This is Part I of a series of two articles reporting the value of in-line XPCT for virtual histology of the brain; Part II shows how in-line XPCT enables the whole-brain 3D morphometric analysis of amyloid-β (Aβ) plaques.

**Highlights:**

- X-ray phase-contrast tomography (XPCT) enables myelin mapping of the whole brain
- XPCT detects and quantifies white-matter injuries in a range of diseases
- Fiber directions and anisotropy metrics can be retrieved from XPCT data
- XPCT is compatible with subsequent conventional histology of brain samples
- XPCT is a powerful virtual histology tool that requires minimal sample preparation

**Graphical Abstract:** 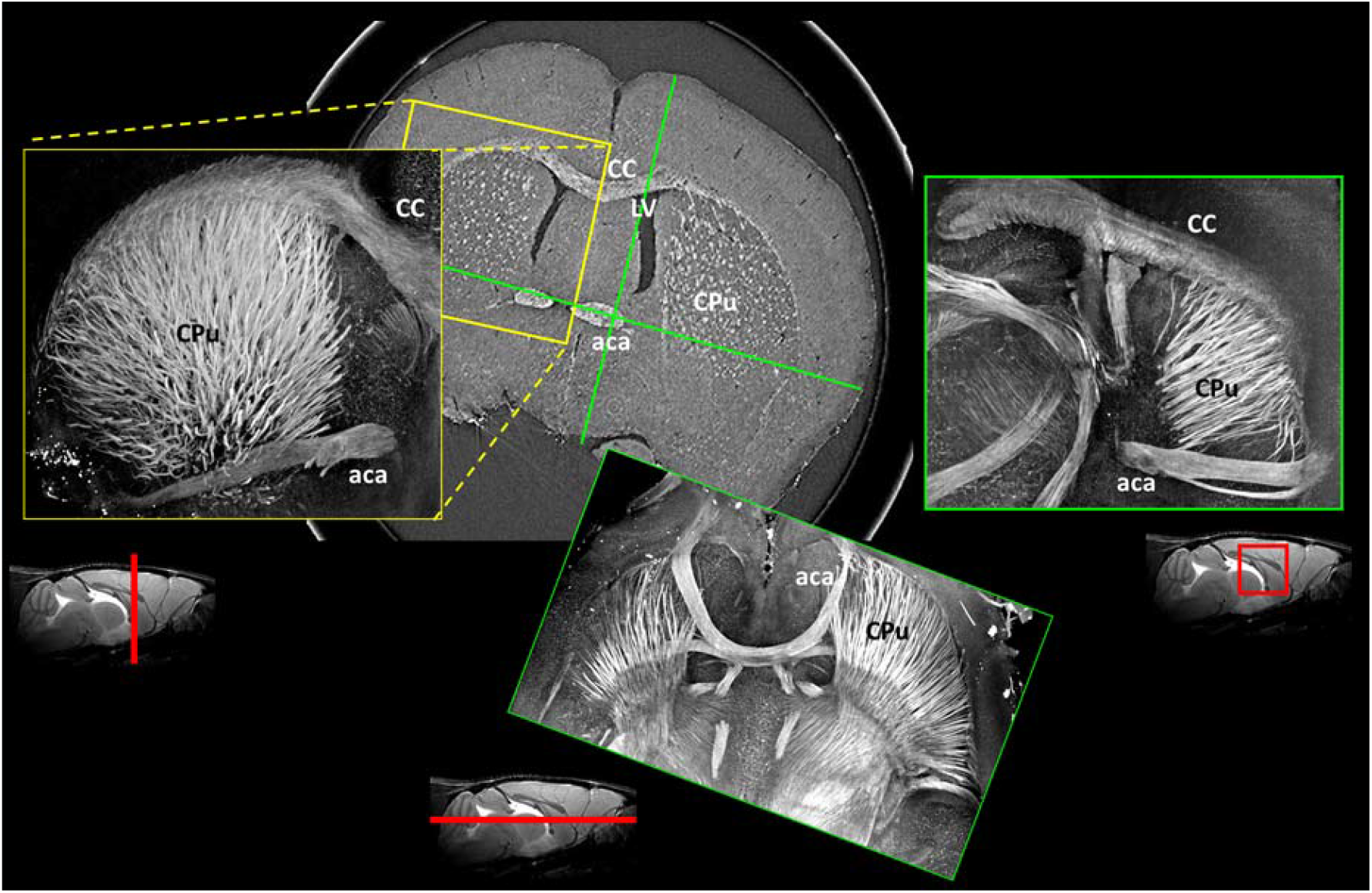

## Introduction

White-matter injury leads to severe functional loss in many neurological diseases. Diagnosis and monitoring of white-matter damage and repair is therefore paramount in order to limit the handicap resulting from these conditions. Magnetic resonance imaging techniques such as diffusion tensor imaging (DTI) and its variants (q-balls, kurtosis imaging) are now instrumental for the assessment of white-matter abnormalities (1). Beside displaying fiber tract orientation maps, these approaches provide quantitative metrics such as fractional anisotropy (FA), axial diffusivity (AD) and radial diffusivity (RD), which in combination serve as biomarkers of axonal and myelin damage (2). However, these techniques still present limitations, notably due to partial volume effects, the difficulty of resolving crossed fibers, and the delicate interpretation of DTI metrics that may be influenced by factors other than axonal degeneration and myelin degradation, such as inflammation and astrogliosis (3). There is thus a crucial need to develop novel whole-brain imaging approaches to study white-matter tracts at the microscopic level (4), especially for the investigation of animal models of white-matter injury.

Myelin staining on histological samples is a fundamental technique employed in many areas of neuroscience to study white-matter fibers. However, tissue processing and sectioning affect the reliability of 3-dimensional (3D) volumetric assessments. Recently, brain-clearing approaches coupled to the use of antibodies, for example targeting myelin basic protein (MBP) to visualize myelinated axons (5), have been proposed to overcome these limitations. Unfortunately, these techniques are laborious and time-consuming. In addition, tissue immunostaining is dependent on antibody penetration into the optically cleared brain, while light-sheet microscopy (the companion imaging tool for imaging cleared tissues) has limited through-plane resolution, thus introducing bias when retrieving 3D information (6).

X-ray based virtual histology is emerging as a new discipline, virtually slicing 3D datasets in any direction at microscopic level (7). X-ray phase-contrast tomography (XPCT) using synchrotron radiation is particularly promising to image microstructures in excised biological tissues that have weak X-ray absorption. XPCT achieves a high signal-to-noise ratio without the need to add staining agents. It offers micrometric spatial resolution and isotropic reconstruction in a ∼cm^3^ field of view (FOV), thus providing the ideal prerequisites for imaging white-matter fibers in the entire and intact (unsliced, unstained) rodent brain. To meet this challenge, however, certain improvements are required, notably to obtain sufficient contrast-to-noise in axons without labor-intensive contrast-enhancing steps (8).

We previously developed an imaging technique relying on in-line XPCT that identifies mouse brain anatomy as clearly as histology, with acquisition time in the range of minutes (9-11). We here propose an optimized approach to map myelin fibers in the whole mouse brain with microscopic resolution without the need for strenuous staining. We provide the proof-of-concept that this approach enhances myelinated fibers in rodent and human brain tissue. In addition, we demonstrated that the technique detected and quantified white-matter injuries using three animal models: ischemic stroke, premature birth and multiple sclerosis. Furthermore, analogously to diffusion tensor imaging (DTI), we retrieved fiber directions and DTI-like diffusion metrics from our XPCT data to quantitatively characterize white-matter microstructure. Finally, we here show that this non-destructive approach is compatible with subsequent complementary brain sample analysis by conventional histology. In-line XPCT might thus become a novel gold-standard for investigating white-matter injury in the intact brain *ex vivo*. This is Part I of a series of two articles reporting the value of in-line XPCT for virtual histology of the brain; Part II shows how in-line XPCT enables whole-brain 3D morphometric analysis of amyloid-β (Aβ) plaques (12).

## Materials and Methods

Table 1 summarizes the samples and imaging techniques used in the study.

**Table 1.** Summary of samples used for XPCT, with concurrent DTI and/or histology.

| Species | Age | Disease / Animal model |  | XPCT | DTI | Histology |
| --- | --- | --- | --- | --- | --- | --- |
| Mouse | 8 weeks | None |  | 7 | 1 | 1 |
|  |  | Ischemic stroke |  | 2 |  | 1 |
|  |  | Multiple sclerosis |  | 2 |  |  |
|  | 11 days | Premature birth | *Hypoxia* | 2 |  | 2 |
|  |  |  | *Normoxia* | 2 |  | 2 |
| Rat | 10 weeks | Multiple sclerosis |  | 1 | 1 |  |
| Human | 70 y.o. | Frontotemporal dementia |  | 2 |  |  |

### Animals

All experimental procedures involving animals and their care were carried out in accordance with European regulations for animal use (EEC Council Directive 2010/63/EU, OJ L 276, Oct. 20, 2010). The study was approved by our local review board “Comité d’éthique pour l’Expérimentation Animale Neurosciences Lyon” (CELYNE - CNREEA number: C2EA – 42, APAFIS#7457-2016110414498389, 5892-2016063014207327, 187-2015011615386357). Sixteen rodents (Janvier, France) were used (Table 1). Animals were housed in a temperature- and humidity-controlled environment (21 ± 3°C), with 12:12h light-dark cycle, with free access to standard chow and tap water. For surgeries, animals were anesthetized with isoflurane (induction: 3.5%; surgery: 2%; ISO-VET, Piramal Healthcare, Morpeth, UK). Pain was alleviated by subcutaneous injection of buprenorphine at 0.05 mg/kg prior to surgery. During surgery, body temperature was monitored with a rectal probe and maintained at 37°C using a feedback-regulated heating pad. At end of surgery, wounds were treated with lidocaine (lidocaine/prilocaine 5%, Pierre Fabre, France).

### Animal models of white-matter injury

Three animal models of white-matter injury were used; the number of animals per model is specified in Table 1. Firstly, focal cerebral ischemia, a model of ischemic stroke encompassing the sensorimotor cortex and adjacent corpus callosum, was induced by permanent occlusion of the distal middle cerebral artery (pMCAO), as previously described (13). Briefly, the right MCA was exposed by subtemporal craniectomy and occluded by electrocoagulating the distal part for 10 minutes. The animals were then allowed to recover from anesthesia and sacrificed at day 9 post-surgery. Secondly, chronic hypoxia was used as a model of very premature birth. This model consists in placing newborn mice in chronic hypoxia (10% O2 for 8 days from post-natal day 3 (P3) to P11) during the early postnatal period, corresponding to the late prenatal period of brain development in humans. This model reproduces all histopathological features observed in very preterm infants, including diffuse white-matter injuries (14). Mice were sacrificed at the end of the hypoxic period: i.e., P11. Thirdly, focal demyelinating lesions, a model of multiple sclerosis, were induced by stereotaxic injection of lysophosphatidylcholine (LPC) (Sigma-Aldrich, ref. L4129) at 1% in saline solution into the corpus callosum as previously described (15)(16). After bilateral craniotomy, LPC and saline were slowly infused (0.1µl/min) with 30-gauge needles via tubing connected to syringes installed in injection pumps. In one mouse, 1-µL was injected at AP 0.0 mm; ML ± 2.5 mm; DV -2.0 mm, with contralateral injection of 1-µL saline. In the other mouse, 2-µL was injected at the same ipsilateral site, without contralateral injection. In the rat, 5-µL was injected: AP -0.3 mm; ML ±3.3 mm; DV -2.8 / -3.5 mm; 2.5-µL each, from depth to superficial, with contralateral injection of 5-µL saline. The animals were then allowed to recover from anesthesia and sacrificed at day 7-9 post-surgery.

### In vivo MRI

For *in vivo* MRI, the animals were placed in a temperature-controlled cradle and anesthetized by isoflurane as for surgery. The respiratory rhythm was monitored by a pressure sensor linked to a monitoring system (ECG Trigger Unit HR V2.0, RAPID Biomedical, Rimpar, Germany). MRI was performed on a horizontal 7T Bruker BioSpec MRI system (Bruker Biospin MRI GmbH, Bruker, Germany) equipped with a set of gradients of 440 mT/m and controlled via a Bruker ParaVision 5.1 workstation. A Bruker birdcage volume coil (inner diameter = 72 mm, outer diameter = 112 mm) was used for transmission, and a Bruker single loop surface coil (15 mm diameter) was used for reception. T2-weighted imaging and DTI were obtained; Suppl. Table 1 summarizes the acquisition parameters.

### Rodent and human brain sample preparation

All animals were euthanized by intracardiac perfusion with phosphate-buffered saline (PBS) and their brains were harvested. For XPCT, the impact of dehydration was examined in healthy brains fixed by formaldehyde 4% and dehydrated in ethanol baths (25%; 50%; 75% and 96%). All subsequent (rodent and human) brains were fixed in formaldehyde 4% and then dehydrated in successive ethanol baths until a titration of 96% was reached. Some of the mouse brains were re-hydrated for histology and immunohistology. In addition to the rodent brains, cortex tissue (2 adjacent blocks of ∼1 cm^3^) from one patient with frontotemporal dementia was imaged; it was obtained from the local brain bank in Lyon: Tissue Tumorothèque Est, CRB-HCL (national authorizations DC2008-72 & AC2015-2576). One healthy mouse and one human brain sample were cleared with the CLARITY system (17) after formaldehyde fixation and before ethanol dehydration: brains were incubated in a polymerization solution composed of PBS buffer 1X, with the addition of VA-044 concentrated at 0.25% and acrylamide 4%. Brains were kept in the solution for 12 hours at 4°C. After infusion, the gel was polymerized at 37°C under gentle agitation for 3 hours. Brains were then placed in the Logos® electrophoretic tissue clearing solution for one day and cleared using the electrophoretic chamber of the X-Clarity system. Brains were cleared at 1.5 A in 6 (mouse) or 8 (human) hours.

### X-ray phase-contrast computed tomography (XPCT)

Imaging was performed at the ID17 and ID19 beamlines of the ESRF, the European Synchrotron (ESRF, Grenoble, France). The samples (whole mouse brains, rat hemispheres) were conditioned in 1 cm diameter plastic tubes filled with ethanol 96%. We used the propagation-based imaging technique, which exploits free-space propagation in order to have detectable interference fringes. An example of the experimental set-up is shown in Suppl. Fig. 1. A summary of experimental and reconstruction parameters is given in Suppl. Table 2. Briefly, the tomographic images were recorded at a single sample-detector distance where the camera was positioned away from the sample to obtain phase contrast. The experiments were performed with a polychromatic “pink” incident X-ray beam (energy: 19-35 keV according to the experiment setting, Suppl. Table 2). Tomographic reconstructions were performed using the Paganin single distance phase-retrieval approach of the PyHST2 software (18). For all scans, the δ/β ratio was set to provide optimal contrast without blurring the image too much (11) (Suppl. Table 2).

**Table 2.** XPCT quantifications. (ROI: region of interest; SD: standard deviation)

| Animal model | Parameter |  |  |  |
| --- | --- | --- | --- | --- |
| Ischemic stroke | Fiber density (%) |  |  |  |
|  | Mouse #1 |  | Mouse #2 |  |
|  | Ipsilateral | Contralateral | Ipsilateral | Contralateral |
|  | 6% | 11% | 11% | 13% |
| Premature birth | Fiber density, mean of left and right ROI ± SD | | | |
|  | Mouse #1 |  | Mouse #2 |  |
| *Normoxia* | 11% ± 2% | | 9% ± 2% | |
| *Hypoxia* | 0% ± 0% | | 0% ± 2% | |
| Multiple sclerosis | Lesion volumes ($10^8 \times \mu\text{m}^3$) | | | |
| | Mouse #1<br>(1-$\mu\text{L}$ LPC) | Mouse #2<br>(2-$\mu\text{L}$ LPC) | Rat<br>(5-$\mu\text{L}$ LPC) | |
|  | 1.2 & 1.8 | 4.5 | 8.7 |  |
|  | DTI-like parameters (% of mirror ROI) |  |  |  |
| | Rat (5-$\mu\text{L}$ LPC) | | | |
|  | FA: 72% | AD: 79% | RD: 146% |  |

### Data processing and quantitative analysis

The mouse brain library was used for labeling white-matter fiber tracks (http://www.mbl.org/mbl_main/atlas.html). Maximum intensity projections were generated using ImageJ software (https://imagej.nih.gov/ij/index.html). Segmentation, analysis and volume rendering were performed with Amira Software (https://thermofisher.com/amira-avizo). For fiber density quantification, XPCT hyperintense signals of the caudate putamen were segmented using a home-made pipeline made with Amira Software. This pipeline relied on a structure-enhancement filter (Frangi filter for rod-like elements (19)), then processed with the TEASAR tree-structure extraction algorithm (20) to segment the white-matter network. Fiber density was then evaluated as the ratio of the number of segmented voxels over the total number of voxels within 3D ROIs. For mice with pMCAO, ROIs encompassing 100 axial slices at the level of the ischemic lesion were placed in the peri-lesional caudate putamen. For mice with chronic hypoxia, ROIs encompassing 60 slices were placed in the center of the caudate putamen. The DTI parametric maps (FA, RD, AD) were reconstructed from MRI data using DSI studio software (http://dsi-studio.labsolver.org). The DTI-like parametric maps were computed from XPCT images using a dedicated tool developed by NOVITOM (https://www.novitom.com/en/). The algorithm was developed based on image gradient analysis to detect fibers orientation. The basic idea is that image gradient should be minimal along the fibers direction. The first step is the computation of the image intensity gradient along the 3 spatial axes, giving a gradient vector for each voxel:

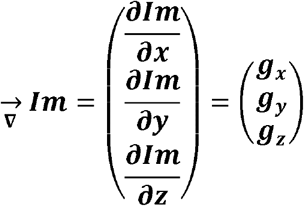

The gradient orientation tensor (or covariance matrix) is then built using the tensor product:

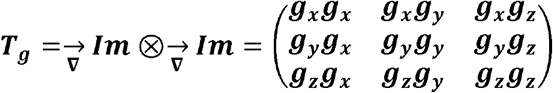

The tensor components are locally correlated using a convolution by a gaussian kernel with a standard deviation ***σ***_***g***_:

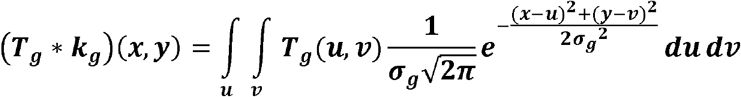

Then the tensor is diagonalized to extract the local orientation axes:

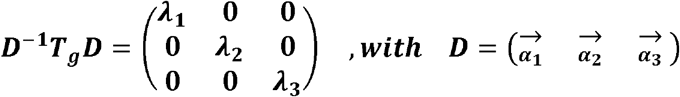

The eigen vector 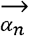 is associated with the eigen values *λ*_*n*_. The eigen vector 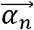 is associated with the smallest eigen value *λ*_*n*_ is aligned with the direction in which the gradient is weakest, which should be locally aligned with the fibers direction. The fibers orientation 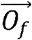 is defined as:

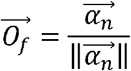

The fibers orientation tensor is constructed as the tensor product of this vector by himself:

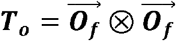

The components of the tensor are locally correlated by applying a convolution by a gaussian kernel with a standard deviation ***σ***_***o***_:

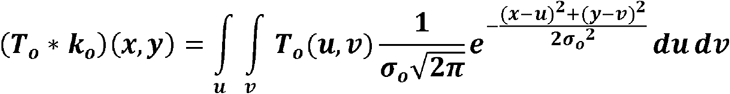

Finally, the orientation tensor is diagonalized to obtained the eigenvalues used to compute the DTI parametric maps (FA, RD, AD).

### Histology and immunohistology

Brain samples were frozen at −80°C. For GFAP and collagen IV staining, rehydrated frozen brains were cut into 12-µm axial sections using a cryostat. Brain slices were rehydrated 10 minutes in 0.1M PBS, fixed for 20 minutes in 4% PFA, and rinsed 3 times with PBS. They were then blocked and permeabilized in PBS (Sigma-Aldrich, Saint Louis, MO, USA) with 5% BSA (bovine serum albumin, Sigma-Aldrich, Saint Louis, MO, USA) and 0.5% Triton X-100 (PBST) for 30 minutes at room temperature. Slides were then incubated overnight at 4°C with primary polyclonal anti-GFAP rabbit antibody (1:1000; Z0334, DAKO, Golstrup, Denmark) or anti-collagen IV rabbit antibody (1:200, ab6586, AbCam, Cambridge, UK) dissolved in 0.5% PBST. They were then washed 3 times in PBS and incubated with a secondary anti-rabbit GFP antibody dissolved in PBST (1:1000, AF546, ThermoFisher, Waltham, MA, USA) for 1 hour at room temperature. Finally, slides were rinsed 3 times in PBS and mounted in aqueous medium containing DAPI (Roti-Mount®, Carl Roth, Karlsruhe, Germany). For Sudan Black B staining, brain slices were post-fixed with 4% formaldehyde in PBS, briefly dehydrated in 70% ethanol and then incubated in 0.1% Sudan Black B solution (Sigma Aldrich, ref. 199664, Saint Louis, MO, USA) at room temperature for 10 min. They were then washed in 70% ethanol for 10–30 minutes and moved into distilled water to be mounted in aqueous medium (Roti-Mount®, Carl Roth, Karlsruhe, Germany). For MBP staining, brains were first rehydrated in 0.1 M PBS, then cut into 50-µm axial sections using a vibratome. Free-floating sections were blocked in TNB buffer (0.1 M PB; 0.05% Casein; 0.25% BSA; 0.25% TopBlock) with 0.4% triton-X (TNB-Tx), then incubated overnight at 4°C with a rabbit anti-MBP primary antibody (1:1000; AbCam, Cambridge, UK), under constant gentle shaking. Following extensive washing in 0.1 M PB with 0.4% triton-X (PB-Tx), sections were incubated with appropriate secondary antibodies conjugated with Alexafluor 488 (1:500; Life Technologies, Carlsbad, Germany) for 4 hours at room temperature. Sections were washed and counterstained with DAPI (300 nM; D1306; Life Technologies, Carlsbad, Germany). Images were acquired using an Axio Scope A.1 fluorescence microscope (4 filters, Carl Zeiss, Oberkochen, Germany) equipped with a x0.63 AxioCam MRc (Carl Zeiss, Oberkochen, Germany).

### Data and Code availability statement

The XPCT raw data cannot be shared at this time due to the large size of the datasets, but can be made available on reasonable request. The DTI-like algorithm is the property of NOVITOM.

## Results

### XPCT enables visualization of myelinated white-matter fibers in the whole mouse brain

We first aimed to evaluate the impact of dehydration on XPCT brain contrast. As reported earlier by ourselves and others (9, 21-23), formaldehyde fixation resulted in faint hypointense contrast of white-matter fiber tracts (Suppl. Fig. 2, 0%). Successive baths of ethanol resulted in appearance of hyperintense signal in myelinated white-matter fibers, with dose effect (Suppl. Fig. 2, 25%-96%). Because ethanol 96% dehydration produced the best visual contrast, we adopted this brain processing technique for the rest of the study. XPCT of dehydrated brain samples provided 3D mapping of all myelinated white-matter tracts, enabling accurate labeling (Fig. 1). Dynamic visualization of virtual slicing (<u>Suppl. movie 1</u>, coronal incidence, and <u>Suppl. movie 2</u>, sagittal incidence) and static visualization of maximum intensity projections (Fig. 1C-D-E-H-I) revealed the complexity of fiber orientations and entanglement in the whole brain (see for example stria medullaris and fasciculus retroflexus in Fig. 1E).

**Figure 1.**
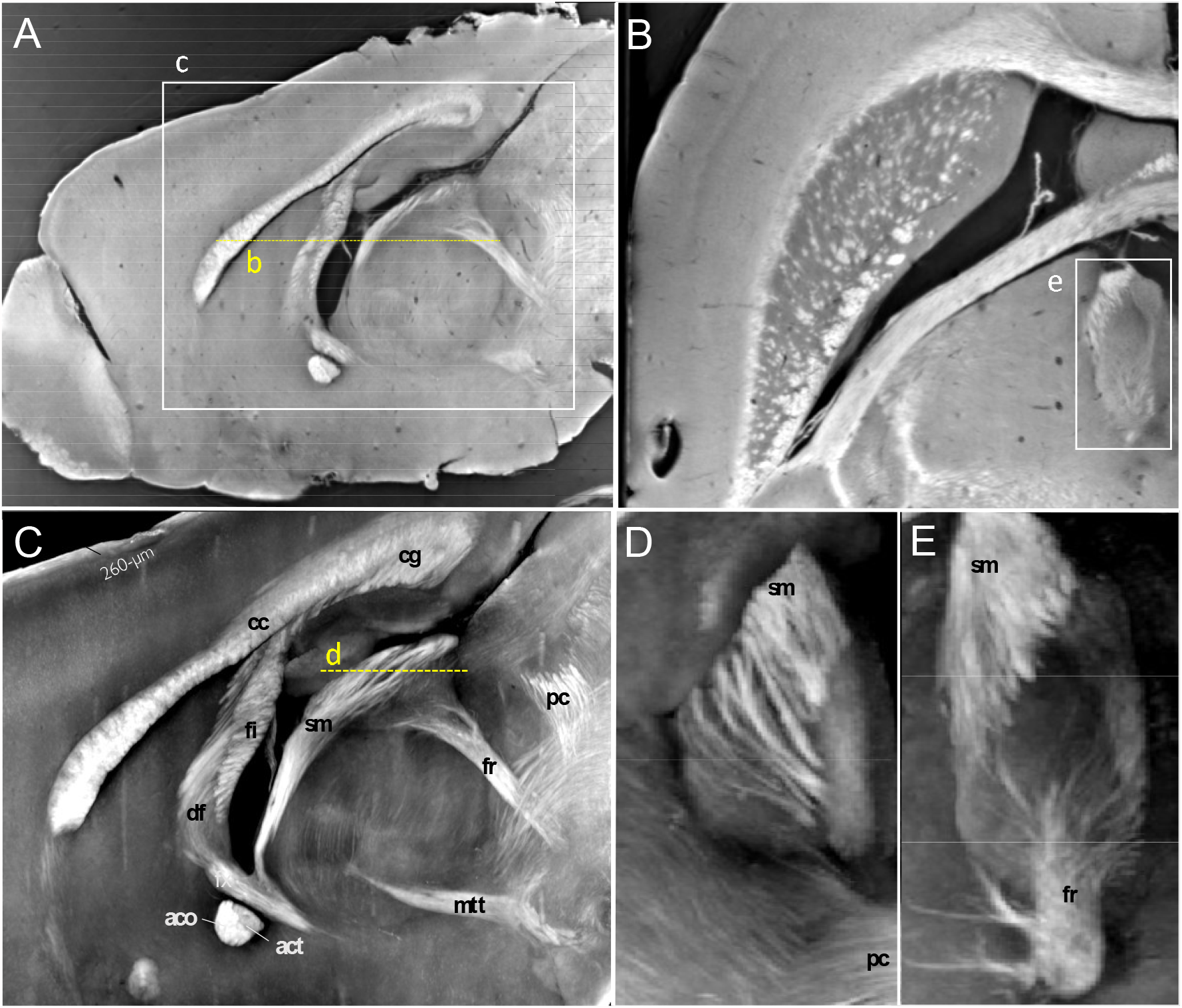
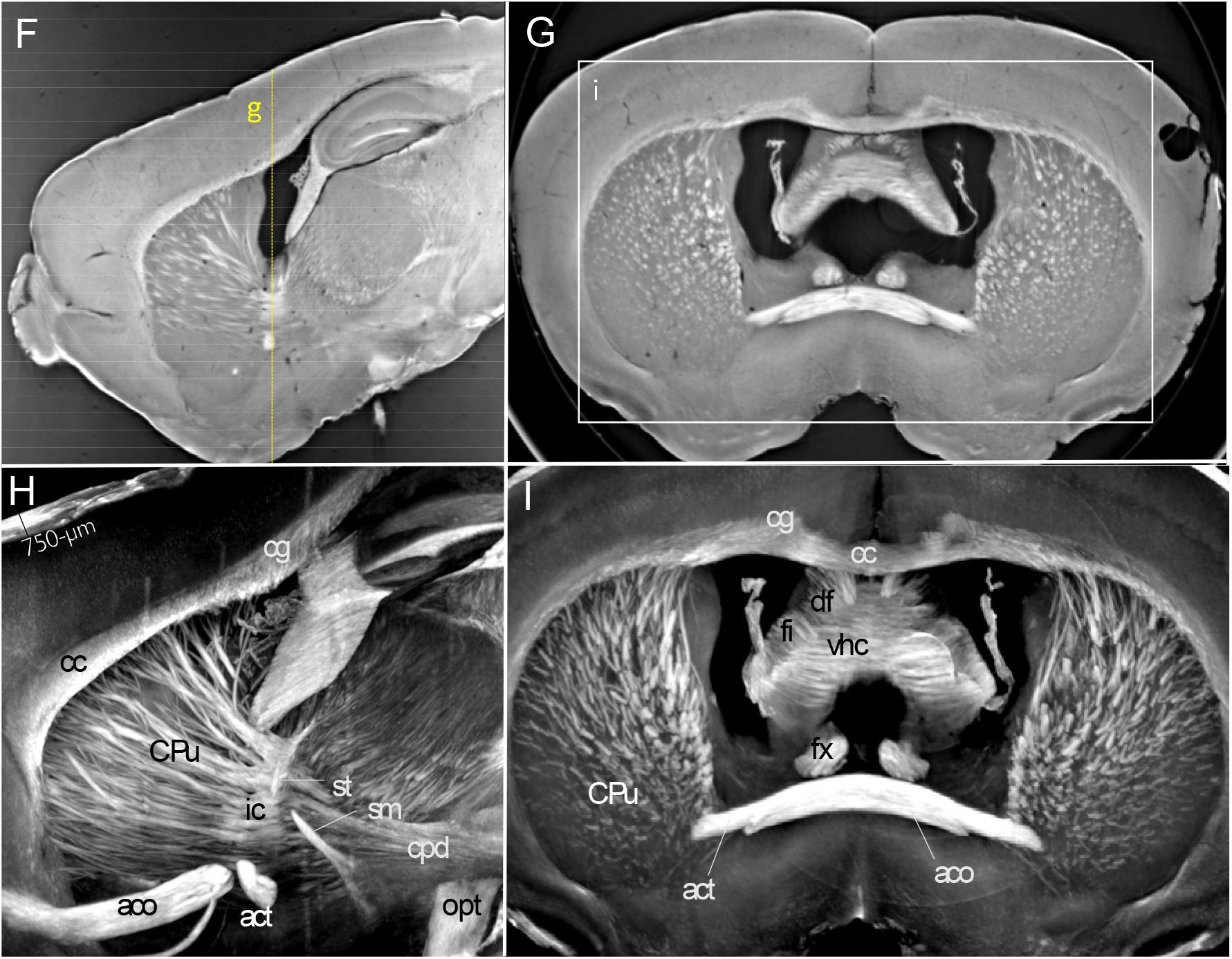
White matter fiber tracts are revealed by XPCT in a mouse brain. (**A**) Sagittal view of single slice (see Movie 1 for 3D animation); (**B**) Corresponding axial view of single slice; (**C**) Maximum intensity projection (MIP) of sagittal view over 260-µm, with white matter fiber tracts annotations; (**D**) MIP showing the entanglement between stria medullaris (sm) and the posterior commissure (pc); (**E**) MIP showing the entanglement between stria medullaris (sm) and the fornix (fr); Labels: aco anterior commissure, olfactory limb; act anterior commissure, temporal limb; cc corpus callosum; cg cingulum; cpd cerebral peduncle; CPu Caudate putamen; df dorsal fornix; fi fimbria; fr fasciculus retroflexus; fx columns of the fornix; mtt mammillothalmic tract; sm stria medularis; st stria terminalis; opt optical tract; pc posterior commissure.; vhc ventral hippocampal commissure. (**F**) Sagittal view of single slice; (**G**) Corresponding coronal view; (**H**) Maximum intensity projection (MIP) of sagittal view over 750-µm, with white matter fiber tract annotations; (**I**) MIP of coronal view over 750-µm, with white matter fiber tract annotations. Labels: aco anterior commissure, olfactory limb; act anterior commissure, temporal limb; cc corpus callosum; cg cingulum; cpd cerebral peduncle; CPu Caudate putamen; df dorsal fornix; fi fimbria; fr fasciculus retroflexus; fx columns of the fornix; mtt mammillothalmic tract; sm stria medularis; st stria terminalis; opt optical tract; pc posterior commissure.; vhc ventral hippocampal commissure.

Importantly, XPCT contrast enhancement of white matter was also observed in human brain tissue after dehydration (Fig. 2A1, arrows).

**Figure 2.**
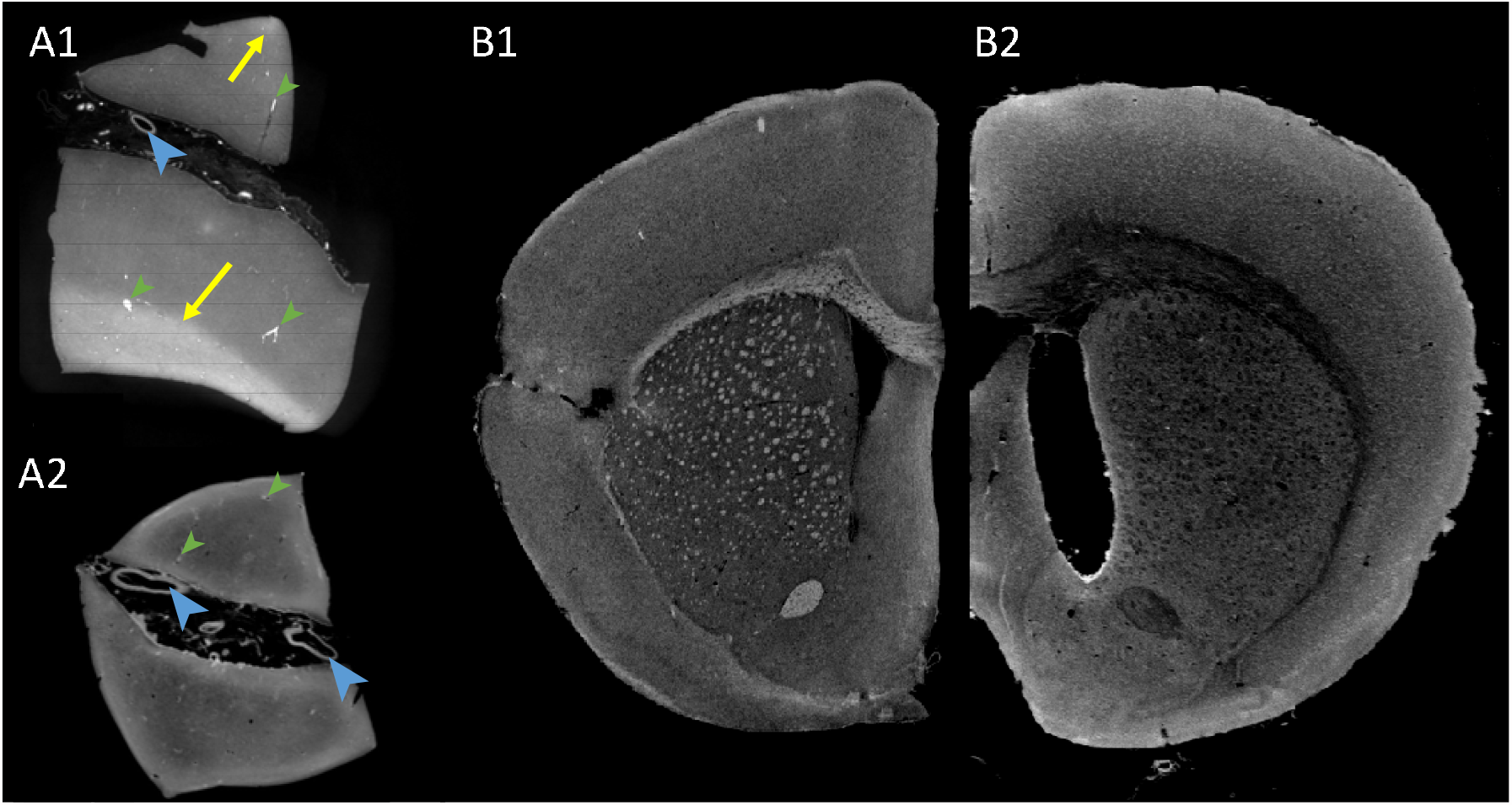
White matter fiber tracts are revealed by XPCT in a human brain and disappear upon demyelination brain clearing. (**A**) Single XPCT slice of a human brain sample (∼1 cm3) from 2 distinct but adjacent blocks. (A1) Ethanol dehydration reveals white matter as hyperintense areas (arrows); (A2) White matter hyperintense signal is lost after demyelination by brain clearing. Other anatomical landmarks such as large vessels are clearly seen (large blue arrowheads). Because the sample was not perfused, there is also contrast from small vessels (smaller green arrowheads); (**B**) Single XPCT slice through a rodent brain without (B1) and with (B2) demyelination by brain clearing: note the disappearance of white matter hyperintense signal.

Brain anatomy was exquisitely revealed by XPCT, with many relevant neurological features (overall vasculature, neuronal bodies of the barrel cortex and hippocampus layers, choroid plexus, cerebellum, etc.) being clearly identifiable (Suppl. Fig. 3).

### XPCT white-matter enhancement in dehydrated brain samples is related to myelin

We hypothesized that the white-matter contrast seen with our approach was related to a mismatch of refractive indices between myelin and surrounding tissue, occurring after the removal of water by ethanol. Myelin has a much higher lipid content than gray matter (24) and might thus be less affected by dehydration than other brain tissues. To investigate this possibility, we used brain clearing, a technique that removes lipids (and hence the myelin sheath) to render the brain translucent to light before ethanol dehydration. We observed a loss of white-matter tract contrast after lipid removal in both human (Fig. 2A2) and mouse brains (Fig. 2B2), suggesting a role of myelin in contrast generation.

### XPCT detects and quantifies white-matter injuries in a range of animal models

We then aimed to evaluate the value of XPCT as a virtual histology tool for the diagnosis and characterization of white-matter injuries. For this, we used 3 different small animal models: i) ischemic stroke (pMCAO), ii) premature birth (neonatal chronic hypoxia) and iii) multiple sclerosis (intracerebral LPC administration).

Firstly, the pMCAO model induced a focal lesion in the sensorimotor cortex, as seen on the in vivo ADC map (Fig. 3A, white dotted lines) and T2-weighted MRI (Fig. 3B, white dotted lines). Subcortical white-matter loss, as can be seen in Fig. 3C (yellow dashed lines), may occur in this model; it is not detectable on conventional MRI and difficult to quantify on histology; with XPCT, 3D segmentation of white matter in the caudate putamen and subsequent quantification of fiber density enabled white-matter loss to be assessed in one individual (Fig. 3D) and white-matter integrity in the other (Table 2).

**Figure 3.**
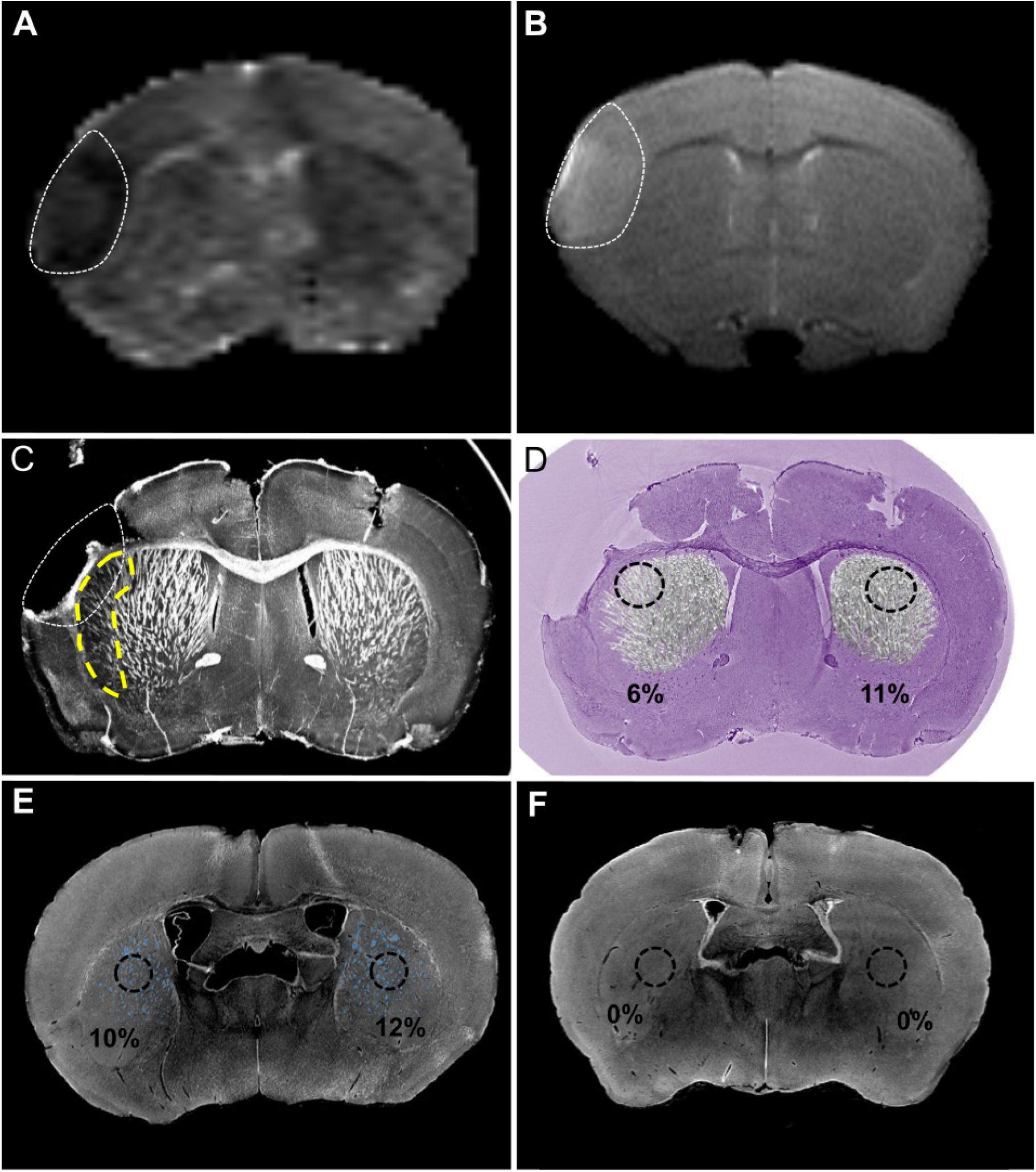
XPCT identifies subcortical white matter loss in mouse models of ischemic stroke and preterm birth. (**A-D**) MRI and XPCT data in a mouse model of ischemic stroke. (A) In vivo MRI data of mouse brain with pMCAO (single slice with 800-µm thickness) imaged at 6 hours post-ischemia. The ischemic lesion (white dotted line) appears hypointense on apparent diffusion coefficient map (A) and hyperintense on T2-weighted image (B); (C) MIP over 800-µm of XPCT data obtained at the same slice level in the same mouse after sacrifice at day 9 post-ischemia. At this stage, the infarcted cortex tended to come off along the perfusion, extraction and dehydration steps of the brain. White matter loss is clearly seen in the subcortical peri-lesional area in this animal (yellow dashed line); (D) Segmentation of caudate putamen white-matter fibers and quantification of fiber density in symmetrical regions (percentage calculated in 3D ROIs over 100 slices). The background XPCT native image is represented in false colors to provide a cresyl violet like contrast. (**E-F**) XPCT data in a mouse model of preterm birth: native (single axial slice) XPCT images in normoxic (E) and hypoxic (F) P11 mice. White-matter fiber tracts were segmented and quantified in the striatum as shown in normoxic animal (E, in blue) but barely detectable in the animals that underwent neonatal chronic hypoxia (F).

Secondly, normoxic mouse neonates showed normal white matter aspect (Fig. 3E) while their hypoxic littermates exhibited very faint white-matter contrast (Fig. 3F). This difference in white-matter content was quantified in the striatum on an individual basis (Table 2).

Thirdly, areas of focal demyelination were clearly observed in LPC-injected animals (Fig. 4A-B, arrow: internal capsule and caudate putamen). Focal lesions remote from the injection site were also seen in one animal, including collapse of the ipsilateral lateral ventricle (Fig. 4A, arrowhead) and demyelination of the olfactory limb of the anterior commissure (Fig. 4C, arrowhead). Disorganization of the stria terminalis was also observed in this mouse, despite the very small size of this fiber tract (Fig. 4D, dashed lines). Segmentation of white-matter abnormalities gave ready access to the volume of demyelinated areas in the individual animals (Fig. 4E, Table 2).

**Figure 4.**
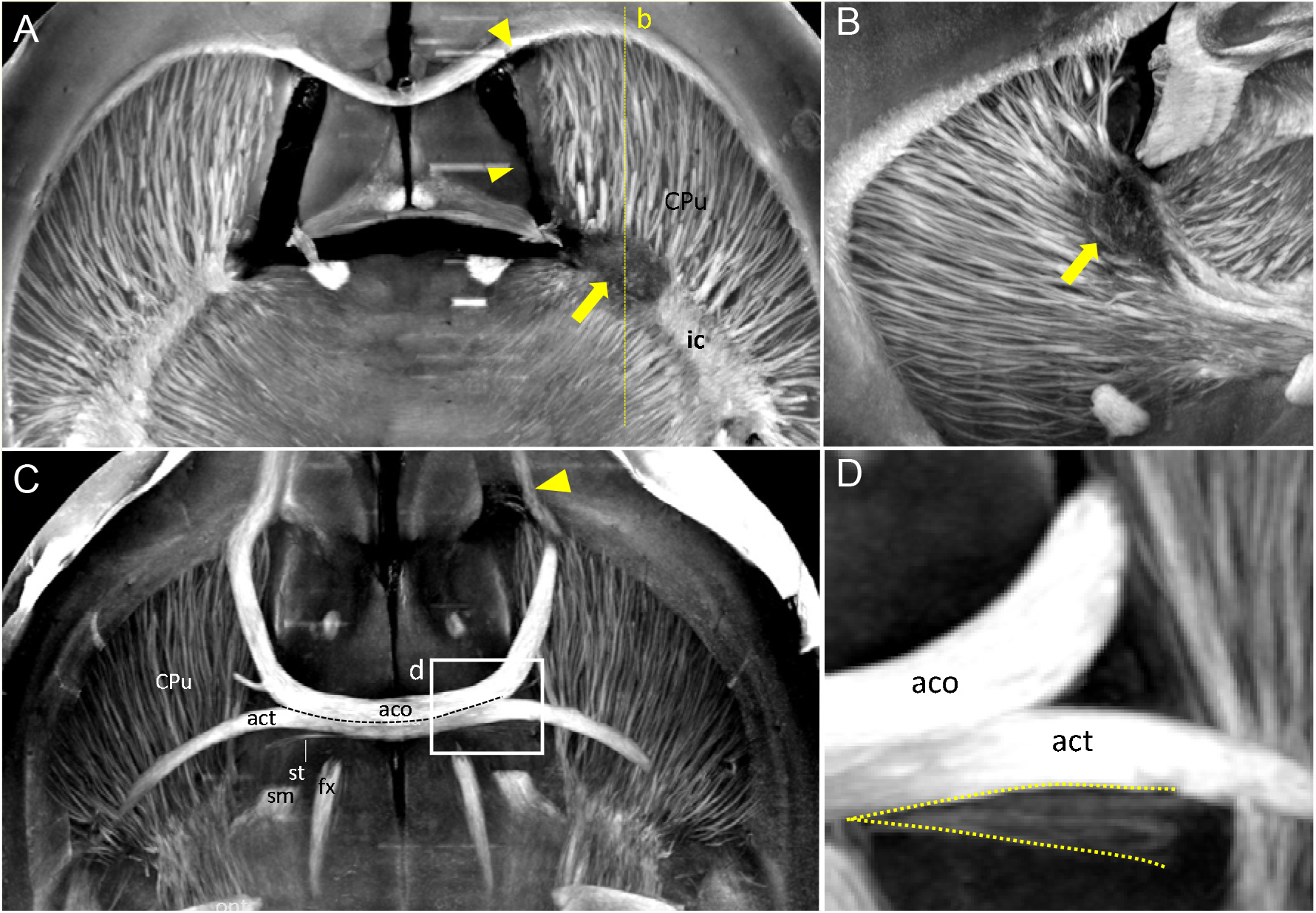
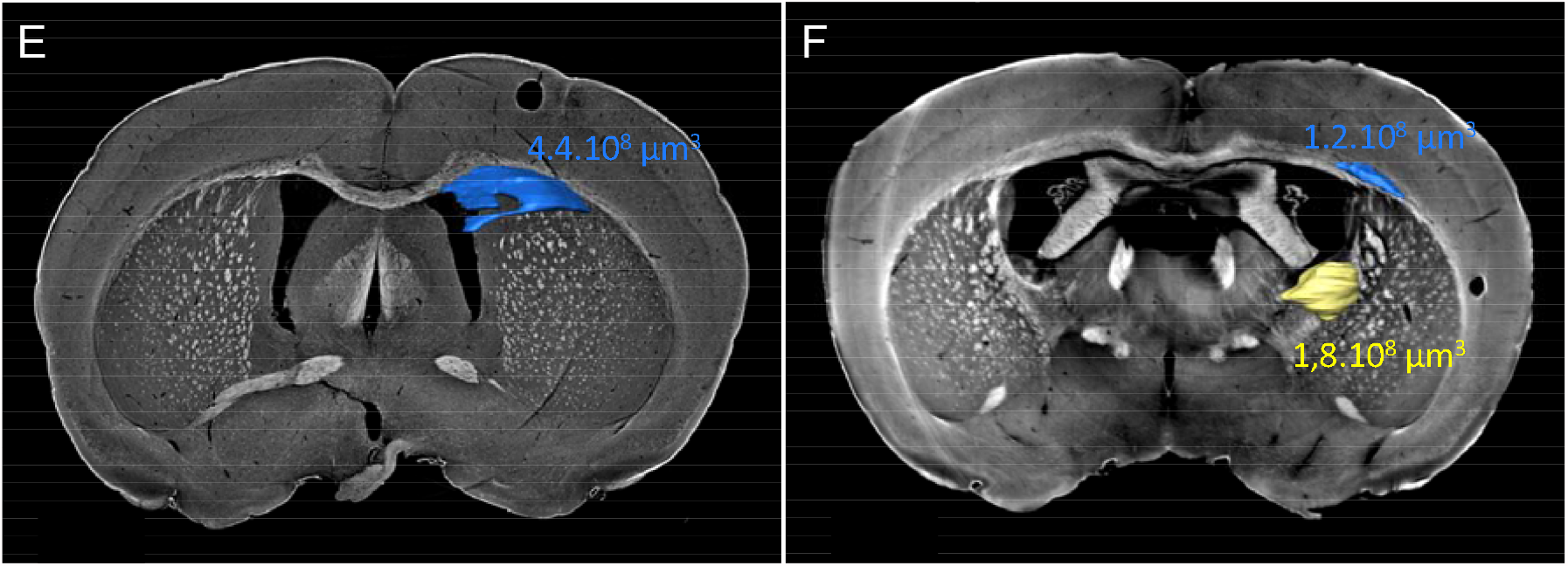
XPCT detects white-matter fiber tract damage in a mouse model of focal demyelination. (**A**) MIP of axial views over 750-µm at the level of the lateral ventricles. Focal demyelination is clearly seen in the internal capsule (arrow). Other focal lesions are shown in the ipsilateral (right) side with an arrow head such as lateral ventricle collapse; (**B**) MIP of sagittal views at the level of the lateral ventricle over 750-µm; Again, focal demyelination is clearly seen in the internal capsule (arrow); (**C**) MIP of axial views over 750-µm at the level of the anterior commissure; degeneration of white-matter fiber tracts can be seen in the ipsilateral anterior part of the anterior commissure (arrowhead); (**D**) Disorganization of the ipsilateral stria terminalis is also seen compared to the contralateral stria, despite the small size of this fiber tract (dashed lines). (**E-F**) Segmentation and quantification of lesion volumes in two separate mice where demyelinated lesions were induced with the LPC model.

### Fiber directions and diffusion metrics can be retrieved from XPCT data

To go further in the analysis of XPCT data, we explored reconstructing the orientation of brain fibers to shed light on brain structural connectivity. A DTI-like algorithm was developed, based on digital gradient analysis to detect fiber orientation. Some of the brain samples (1 normal mouse, and 1 LPC-injected rat) were imaged back-to-back with in vivo DTI (Table 1). Visual examination of brain-wide connectivity maps showed good agreement between the 2 techniques (Fig. 5A-C). LPC-induced demyelination of the corpus callosum was clearly depicted on XPCT fiber-orientation maps (Fig. 5D-E), whereas on DTI maps it was less clear (Fig. 5F). Fig. 5G-H shows examples of diffusion metrics maps obtained with in vivo DTI and XPCT respectively in the same animal.

**Figure 5.**
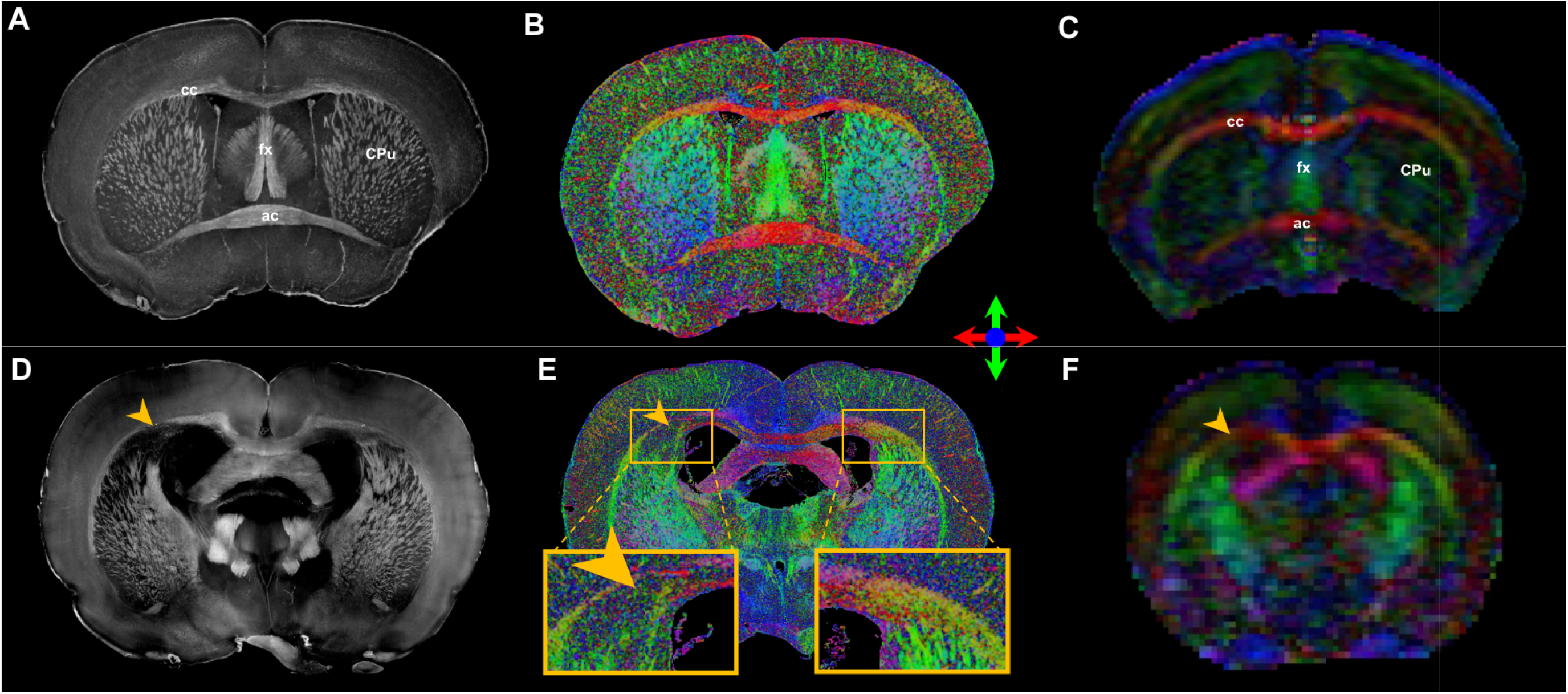
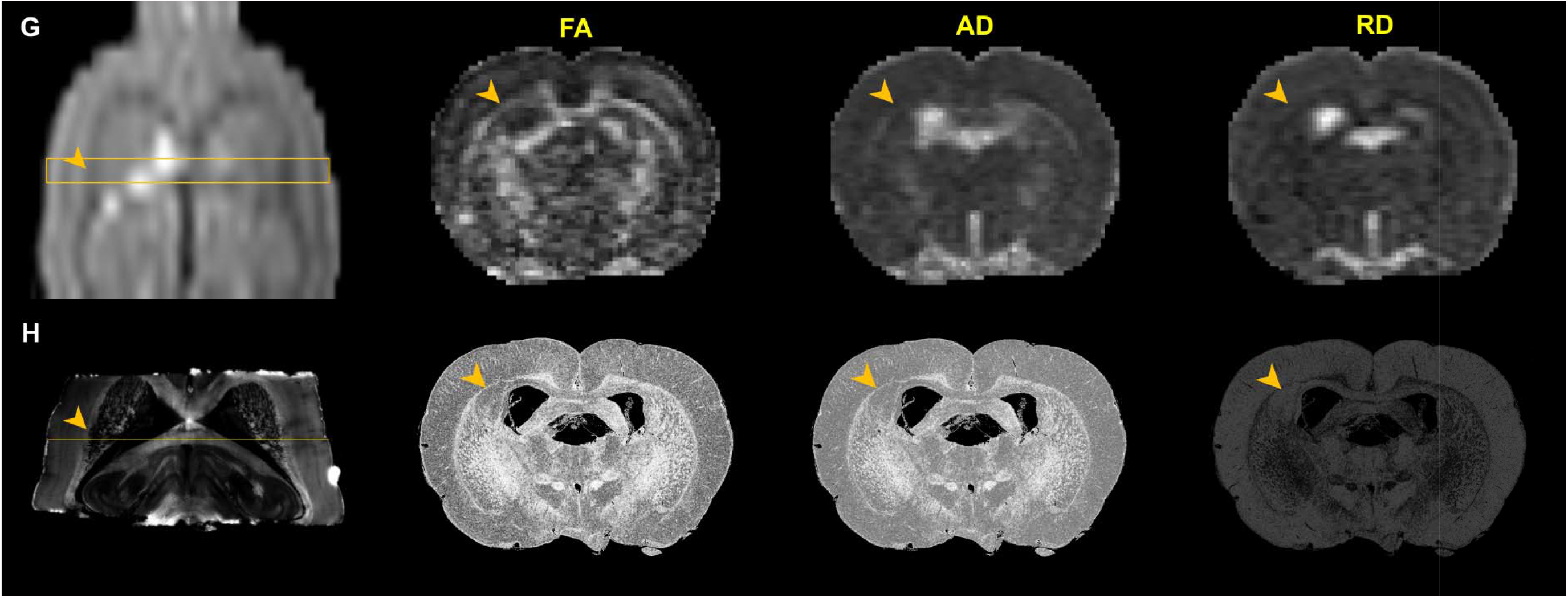
XPCT retrieves brain-wide structural connectivity in a DTI-like manner. (**A**) MIP over 33-µm of a healthy mouse brain (coronal view); (**B**) Color-coded direction map extracted from XPCT from the same data set; (**C**) Color-coded direction map from in vivo DTI obtained in the same mouse at the corresponding slice level. White-matter tract directions are identical between the two techniques within major white-matter tracts such as corpus callosum (cc), fornix (fx), anterior commissure (ac) and caudate putamen (CPu); (**D**) MIP over 65 µm of a rat brain with an LPC-induced lesion in the corpus callosum (coronal view, arrowhead); (**E**) Color-coded direction map extracted from XPCT from the same dataset. The loss in right-left directionality is clearly seen in the demyelinated area compared to the contralateral side (inserts); (**F**) Color-coded direction map from in vivo DTI obtained in the same rat at the corresponding slice level. (**G**) Native DTI data (b0) for a rat with an LPC lesion (coronal view). The box indicates DTI slice thickness (1000-µm) and the arrow points to the area of demyelination. DTI metrics maps of this slice are shown in axial incidence. FA: Fractional anisotropy; AD: Axial diffusivity; RD: Radial diffusivity. (**H**) MIP over 65-µm of XPCT image and DTI-like metrics maps corresponding to the slice indicated on the MIP (slice thickness: 6.5-µm).

Quantification of these metrics in the injured corpus callosum further confirmed the ability of XPCT to quantitatively assess microstructural white-matter abnormalities, with a marked increase in DTI-like RD and a smaller decrease in DTI-like FA and AD compared to the symmetrically contralateral region (Table 2), as expected in demyelinated areas with limited axonal damage (2).

### XPCT is compatible with subsequent conventional histology of brain samples

Finally, we aimed to ascertain that brain imaging with XPCT did not prevent complementary analysis with on standard approaches such as histological staining and immunohistochemistry. After XPCT imaging, 3 brain samples were rehydrated and processed using our conventional 2D histology protocols. GFAP (Fig. 6A), collagen type IV (Fig. 6B), Sudan Black B (Fig. 6C) and myelin basic protein (MBP) (Fig. 6D) immunostaining was readily achievable, with no noticeable impact of prior processing and imaging on staining quality.

**Figure 6.**
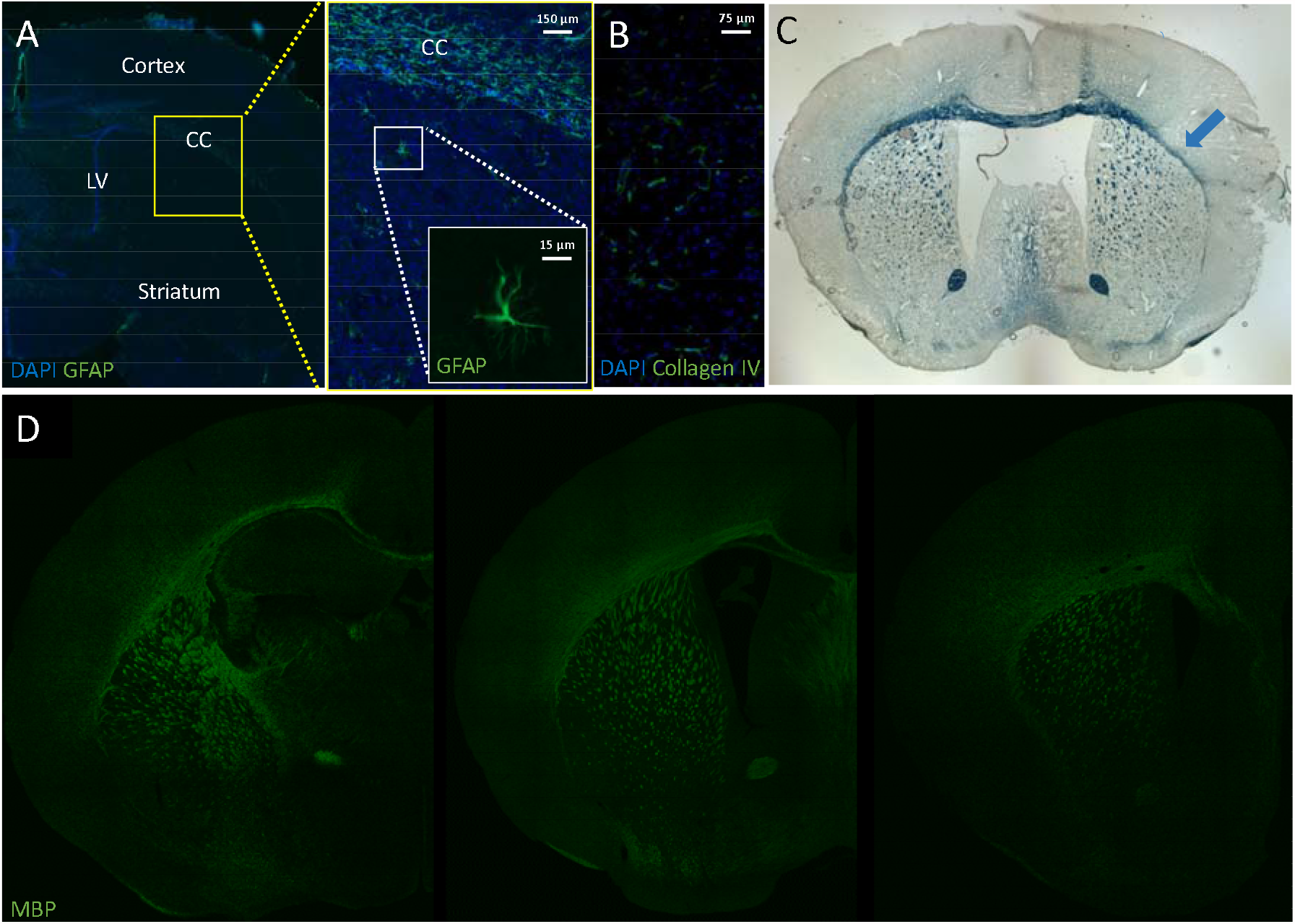
Brain preparation for XPCT does not prevent further brain evaluation with immunohistochemistry. (**A**) Constitutive expression of astrocytes GFAP in healthy mouse (CC: corpus callosum, LV: lateral ventricle); (**B**) Collagen IV overexpression in the ischemic lesion of a pMCAO mouse; (**C**) Sudan Black B staining (myelin marker) of a pMCAO mouse with demyelination of the corpus callosum (blue arrow) and (**D**) Myelin basic protein (MBP) staining in neonatal mice (P11); three slice levels are shown.

## Discussion

This paper describes a novel label-free method based on in-line XPCT for mapping myelin in the whole mouse brain. XPCT offers many advantages over other intact-brain approaches such as electron microscopy (25, 26), brain clearing (5), small angle X-ray scattering (SAXS) CT (6, 27) or dark field CT (28). Firstly, the FOV-to-resolution ratio is optimal for imaging the whole rodent brain at microscopic level. Secondly, isotropic resolution enables straightforward analysis of the 3D imaging data. Thirdly, acquisition and reconstruction times remain within a range of minutes. And last but not least, phase sensitivity offers an alternative to staining, as contrast between tissues is based on changes in X-ray phase rather than on absorption. Consequently, sample preparation is simple and fast. Taken together, this should enable high-throughput whole-brain studies in small laboratory animals and 3D evaluation of human brain samples.

However, phase contrast alone is not sufficient to enhance white-matter tracts, probably because the differences in refractive index between axons and surrounding tissue are minimal. The originality of our approach was to obtain myelin specificity simply by modulating the refractive indices through dehydration of brain samples with ethanol. Ethanol fixation has already been used for virtual XPCT histology of whole organs: vocal folds (29), lung (30), heart (30, 31), and kidney (30, 32, 33). A few brain studies also mentioned using ethanol dehydration to prepare their sample; however, they focused on displaying the angio- and/or cyto-architecture of the brain, or neuropathological features such as amyloid plaques, and did not report specific enhancement of myelinated axons (22, 23, 34-36). Therefore, to the best of our knowledge, we are the first to specifically investigate and demonstrate the value of combining in-line XPCT with ethanol dehydration for 3D visualization of white-matter fiber tracts in the whole mouse brain, to relate the enhanced signal to myelin, and to provide a comprehensive framework for applications in neurological diseases.

Beside its well-known vital role in conducting signals along the nerves, myelin has many other indispensable functions in the central nervous system: providing metabolic support to axons, regulating ion and water homeostasis, and finely tuning neuronal signaling (37). Consequently, disruption of myelin has significant neurological impact. Investigation of myelin disruption and loss in animal models is relevant not only for pathophysiology but also for lesion repair and drug discovery. We here showed, in three different animal models, that XPCT is a powerful tool to probe white-matter injury at organ scale. 3D regions of myelin loss were accurately quantified, and lesions remote from the primary injury site were straightforwardly identified. These intact-brain analyses are a major improvement over classical histology, which involves slicing the brain tissue: i.e., disrupting the structural integrity of axonal tracts.

The technique of directional diffusion is widely used to study white-matter microstructural connections (38). However, DTI has limited in-plane and through-plane spatial resolution, and provides indirect observation of white-matter fibers. Moreover, even when performed *ex vivo* at high resolution, partial volume effects may impair DTI metrics accuracy, and the very long scanning time (typically overnight) is prone to vibration artefacts and tissue heating. We here show that XPCT can be used to obtain an estimate of whole-brain axonal tract orientations in microscopic voxels. This information can then be combined with a directional algorithm to reconstruct structural connections. Classical diffusion metrics (fractional anisotropy, axial, and radial diffusivities) can also be extracted from the diffusion tensor obtained with this algorithm. Thus, XPCT can detect subtle changes in white-matter microstructure in the course of a neuropathology, as illustrated by our results in the LPC rat model of focal demyelination. This method also seems ideally suited for future phenotype screening of myelin abnormalities in transgenic animals. Another potential application of connectome mapping by XPCT is the need for new imaging tools to cross-validate DTI (39). In particular, XPCT might provide a reference in regions of crossing fibers, which are typically difficult to resolve on DTI. Additionally, interpretation of DTI metrics in terms of neuronal and non-neuronal cell types might benefit from XPCT solely depiction of myelinated axons.

Another advantage is that XPCT is non-destructive, and dehydration by ethanol is reversible; thus, brain samples remain available for further analysis. After rehydration, we were able to perform our usual histological and immunohistological analyses. Importantly, brain slices did not show overt signs of radiation damage. In addition, ethanol dehydration/rehydration induces less volume variation and brain sample distortion than brain-clearing techniques. Because XPCT gives access to anatomic landmarks, co-registration of X-ray and optical images may be readily obtained. For instance (relevant to our discussion of DTI validation), affine or non-linear co-registration between DTI, XPCT and brain slices immunolabeled for activated glial cells could help decipher the contribution of inflammation to DTI metrics. Ultimately, such pipelines would pave the way to atlas-based analyses of XPCT datasets (40). Like any technique, XPCT is not devoid of limitations. Imaging live brains at such high spatial resolution is not currently an option, because of dosimetry and because the skull interferes with X-ray coherence. Nevertheless, there is no obstacle to imaging postmortem human brains (6, 22), which has important implications for anatomopathological studies (35, 41). In addition, a synchrotron X-ray source is currently needed to produce high coherence for phase-contrast imaging, which may be a concern in terms of accessibility. However, producing phase contrast with laboratory and clinical sources is an active field of research that might solve the problem in the near future (42, 43). Finally, XPCT produces large datasets that are not easy to handle and necessitate the development of automatic post-processing tools and pipelines. This was achieved in the current project through a public-private partnership. In part II of this series of articles, we develop and share a pipeline designed with open-source tools (https://zenodo.org/record/4584753) to segment amyloid-β plaques and extract 3D morphometric parameters in different animal models of Alzheimer’s disease (12). The development of such end-user-friendly workflows is crucial to allow widespread use of XPCT in the neuroscience community. Collaborative efforts by multidisciplinary teams will be crucial to boosting innovation in the field of novel X-ray imaging techniques and to fostering discoveries that will ultimately benefit patients with white-matter injuries.

## Conclusion

The proposed XPCT approach enables brain-wide studies of myelinated fiber tracts and of their microstructural changes in neurological diseases. This is achieved with conventional animals (i.e., no need for fluorescent reporter mice), minimal sample preparation, and fast acquisition. In the future, combining different imaging techniques with different resolution levels and complementary information will push the frontiers of our understanding of brain structure and function.

## Acknowledgments

This work was performed within the framework of LABEX PRIMES (ANR-11-LABX-0063) of Université de Lyon, within the “Investissements d’Avenir” program (ANR-11-IDEX-0007) operated by the French National Research Agency (ANR). The MRI part of the study was performed on CERMEP imaging facilities (www.cermep.fr, Lyon, France). The authors would like to thank Jean-Baptiste Langlois from CERMEP for animal preparation and MRI acquisition and Corinne Perrin from the Tumorothèque Est tissue bank, CRB-HCL (Lyon, France), for managing human brain samples. We thank Clément Tavakoli for designing Suppl Fig 1. The authors would like to thank our local ESRF contacts Lukas Helfen and Vincent Fernandez, as well as Max Langer, Loriane Weber and David Rousseau for participating in XPCT data acquisition at ESRF. This work was supported by the ESRF by allocation of beam time (LS2292, MD1018, MD1094, MD1106, MD1125). The research was funded by the French national research agency (ANR) NanoBrain (ANR15-CE18-0026), and Breakthru (ANR-18-CE19-0003). LF is supported by ANR NeoRepair (ANR-17-CE16-0009).

## Supplemental Table and Figures

**Suppl Table 1.**
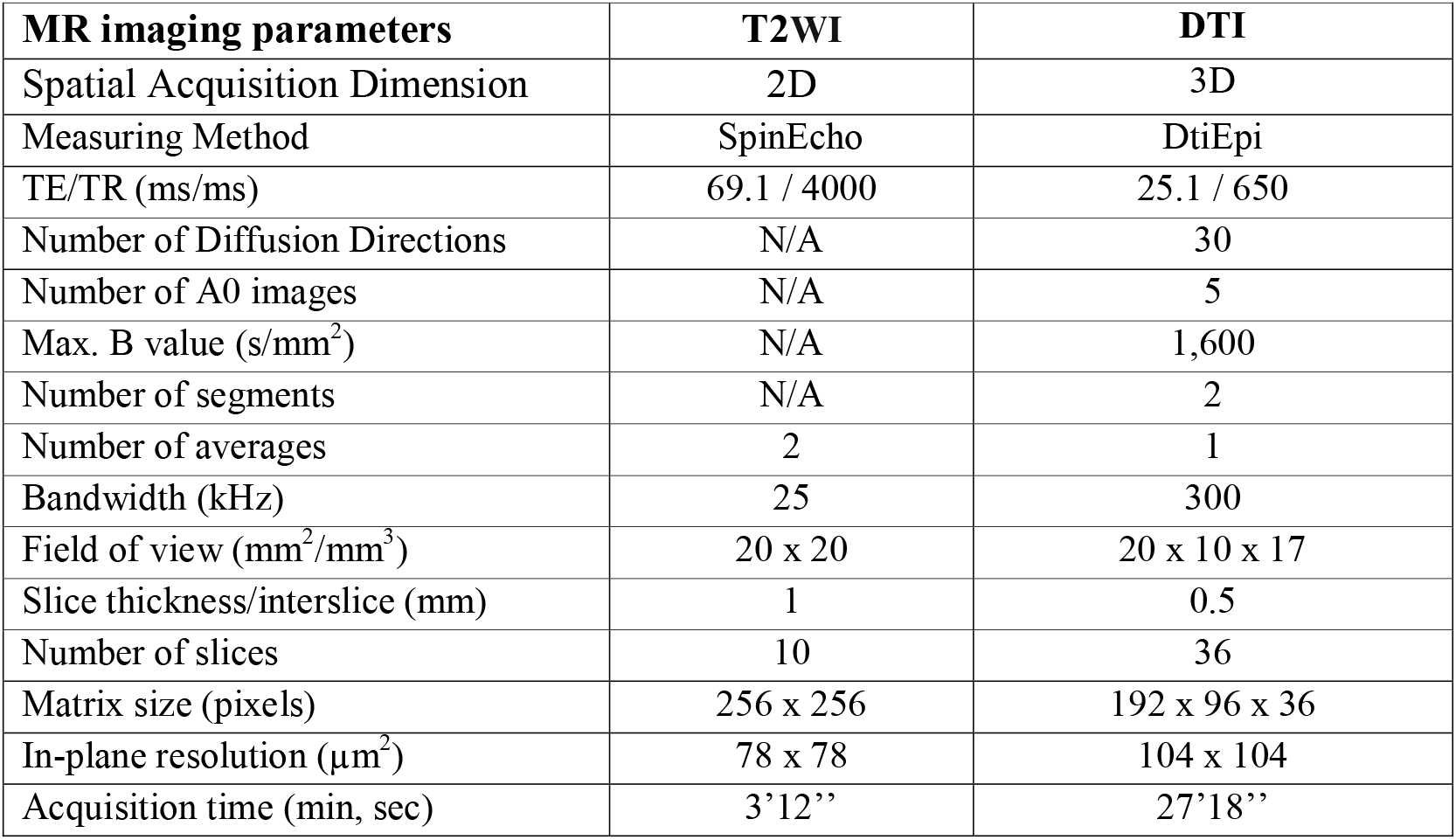
Experimental MRI parameters. (TE: echo time; TR: repetition time; T2WI: T2-weighted MRI; DTI: diffusion tensor imaging; N/A: non-applicable)

**Suppl Table 2.** Experimental XPCT parameters.

| **Figures** | 1, 3C-D, 4, 5A-B, 6B | 2B, 3E-F, 5DE & GH | 2A |
| --- | --- | --- | --- |
| **Supplemental Figures** | 2, 3E-F | 3C-D | 3A |
| **Setup** | ID19 | ID19 | ID17 |
| **Detector** | FReLoN | sCMOS (PCO edge) | sCMOS (PCO edge) |
| **Scintillator** | LuAg 500 μm | LuAg 500 μm | LuAg 500 μm |
| **Energy [keV]** | 19 | 26 | 35 |
| **Sample-detector distance [m]** | 1 | 1.3-3 | 11 |
| **Field-of-view [cm<sup>3</sup>]** | 1.2 x 1.2 x 1.2 | 1.2 x 1.2 x 1.2 | 1.5 x 1.5x 1.5 |
| **Matrix** | 2,048 x 2,048 | 2,048 x 2,048 | 2,560 x 1,000 |
| **Total slice number** | 2,000 | 2,000 | 3,200 |
| **Voxel size (isotropic) [μm]** | 7.5 | 6.5 | 6.13 |
| **Exposure time [sec]** | 0.15 | 0.05 | 0.05 |
| **Number of projections** | 2,000 (180°) | 2,499 (180°) | 2,800 (360°) |
| **Acquisition time [min]** | < 10 min |  |  |
| **Reconstruction algorithm** | Paganin |  |  |
| **δ/β** | 600-1,000 |  |  |

**Suppl Fig 1.**
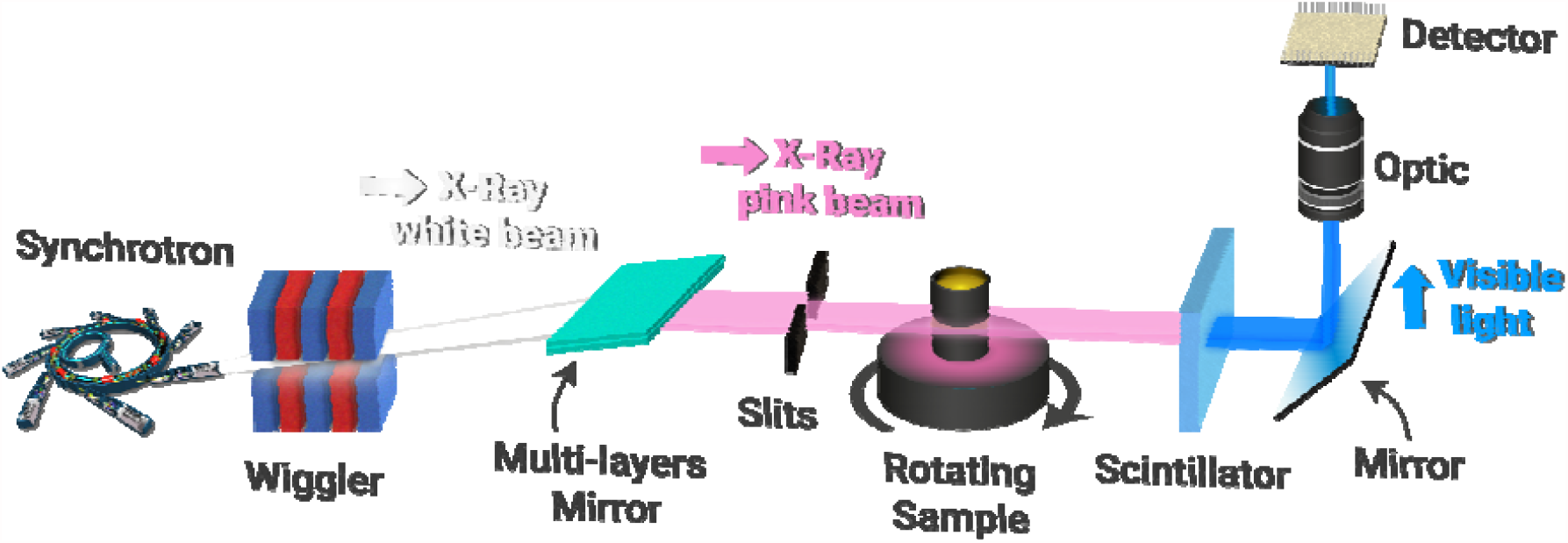
Principle of in-line phase contrast tomography: experimental set-up.

**Suppl Fig 2.**
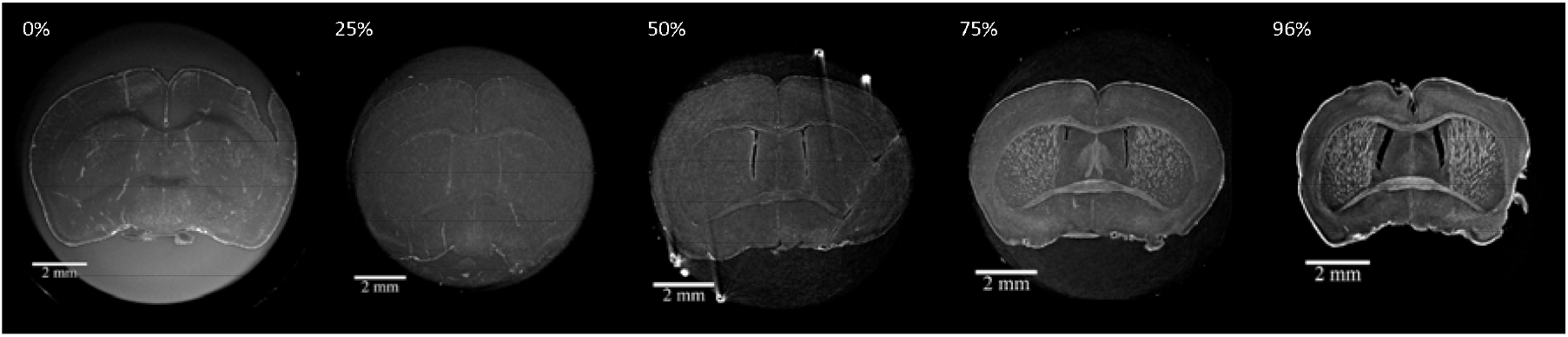
Ethanol dehydration enhances myelin contrast on images obtained with in-line XPCT. Axial section (Bregma 0) through brains prepared using PFA 4% and a gradient of ethanol baths [0%-96%].

**Suppl Fig 3.**
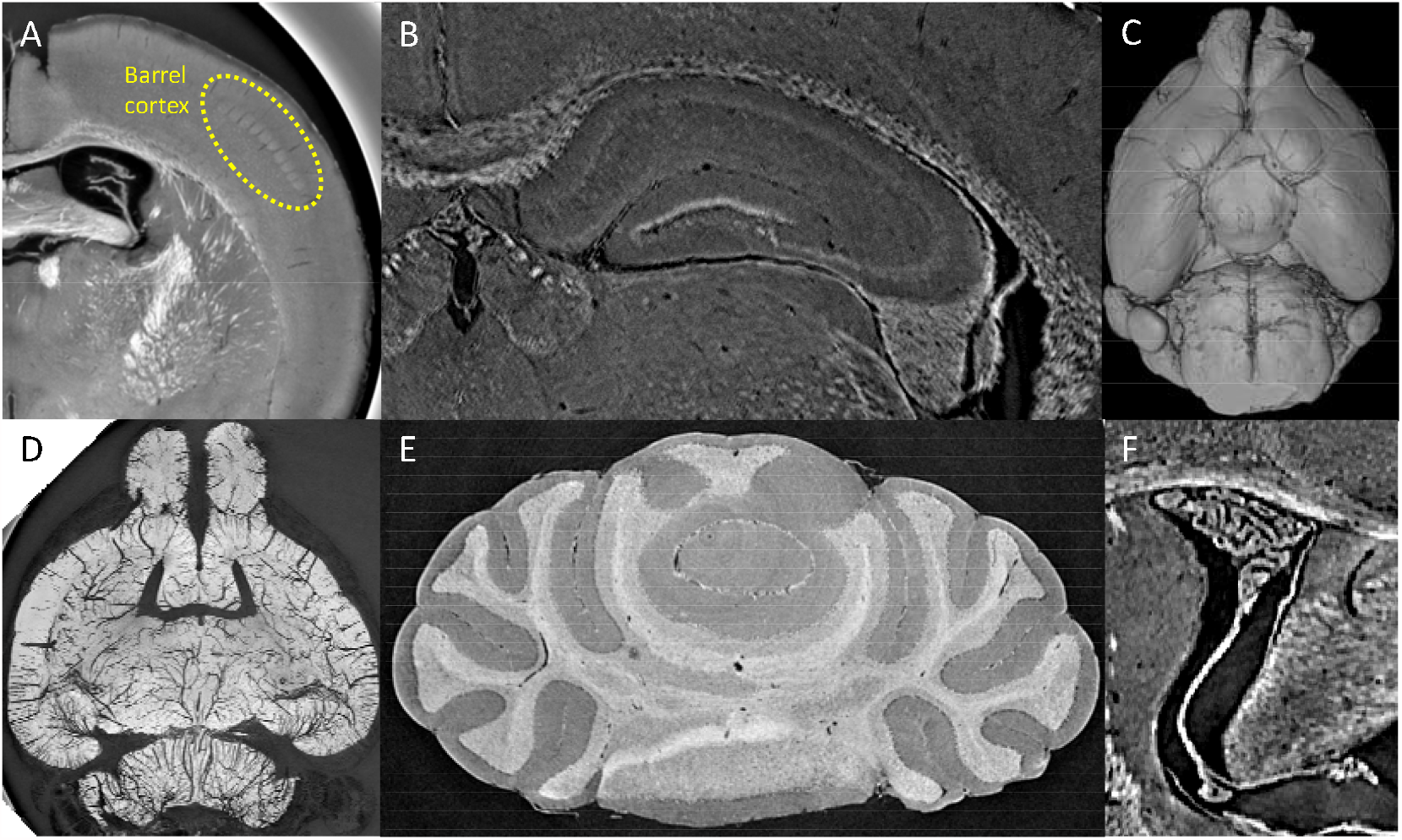
Virtual histology with XPCT displays major brain anatomic features in the whole mouse brain. (A) Barrel cortex; (B) Hippocampus; (C) Circle of Willis; (D) Angiography; (E) Cerebellum; (F) Choroid plexus.

